# Inferring species interactions from co-occurrence data with Markov networks

**DOI:** 10.1101/018861

**Authors:** David J. Harris

## Abstract

Inferring species interactions from co-occurrence data is one of the most controversial tasks in community ecology. One difficulty is that a single pairwise interaction can ripple through an ecological network and produce surprising indirect consequences. For example, the negative correlation between two competing species can be reversed in the presence of a third species that is capable of outcompeting both of them. Here, I apply models from statistical physics, called Markov networks or Markov random fields, that can predict the direct and indirect consequences of any possible species interaction matrix. Interactions in these models can also be estimated from observed co-occurrence rates via maximum likelihood, controlling for indirect effects. Using simulated landscapes with known pairwise interaction strengths, I evaluated Markov networks and six existing approaches. The Markov networks consistently outperformed other methods, correctly isolating direct interactions between species pairs even when indirect interactions or abiotic factors largely overpowered them. Two computationally efficient approximations, based on controlling for indirect effects with linear or generalized linear models, also performed well. Indirect effects reliably caused a common null modeling approach to produce incorrect inferences, however.

## Introduction

To the extent that nontrophic species interactions (such as competition) affect community assembly, ecologists might expect to find signatures of these interactions in species composition data (MacArthur 1958, Diamond 1975). Despite decades of work and several major controversies, however (Lewin 1983, Strong et al. 1984, Connor et al. 2013), existing methods for detecting competition’s effects on community structure are unreliable (Gotelli and Ulrich 2009). In particular, species’ effects on one another can become lost in a web of indirect effects. For example, the competitive interaction between the two shrub species in Figure 1A is obscured by their shared tendency to occur in unshaded areas (Figure 1B). While ecologists have long known that indirect effects can overwhelm direct ones at the landscape level (Levine 1976), the vast majority of our methods for drawing inferences from observational data do not control for these effects (e.g. Diamond 1975, Strong et al. 1984, Gotelli and Ulrich 2009, Veech 2013, Pollock et al. 2014). To the extent that indirect interactions like those in Figure 1 are generally important, existing methods will not provide much evidence regarding species interactions.

**Figure 1:**
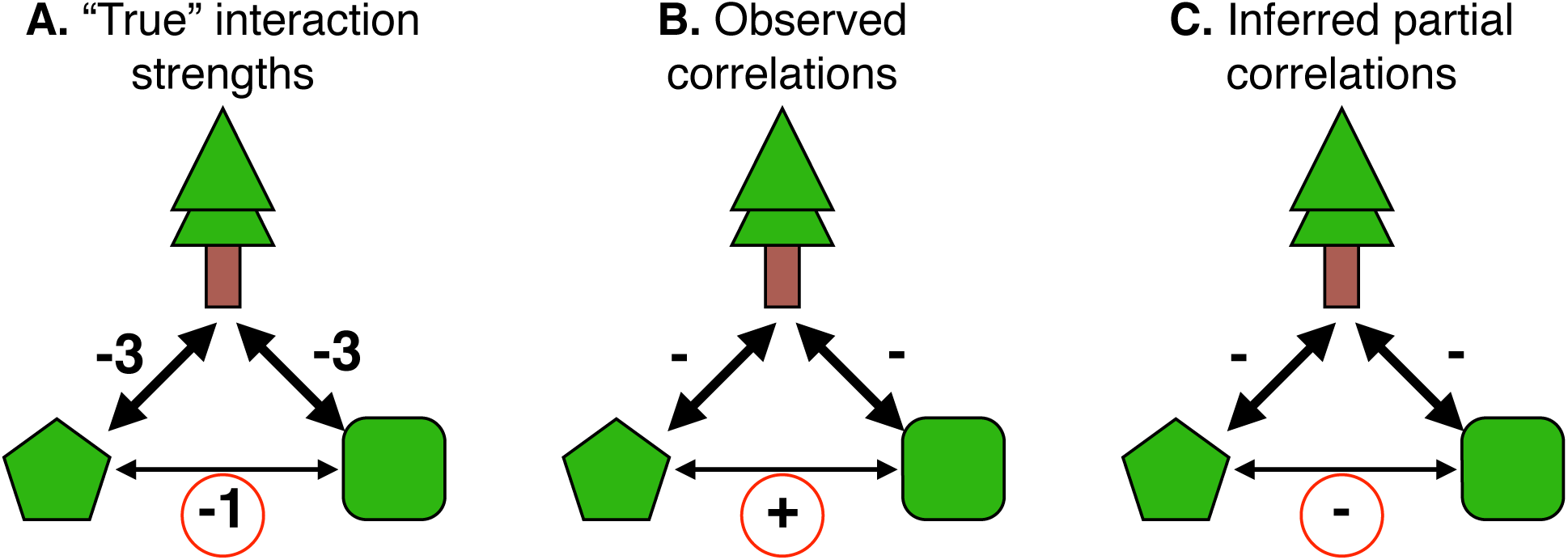
**A.** A small network of three competing species. The tree (top) tends not to co-occur with either of the two shrub species, as indicated by the strongly negative coefficient linking them. The two shrub species also compete with one another, but more weakly (circled coefficient). **B.** In spite of the competitive interactions between the two shrub species, their shared tendency to occur in locations without trees makes their occurrence vectors positively correlated (circled). **C.** Controlling for trees with a conditional (all-else-equal) approach such as a partial covariance or a Markov network leads to correct identification of the negative shrub-shrub interaction (circled).

While competition doesn’t reliably reduce co-occurrence rates at the whole-landscape level (as most methods assume), it does still leave a signal in the data (Figure 1C). For example, after partitioning the data set into shaded and unshaded sites, there will be co-occurrence deficits in each subset that wouldn’t otherwise be apparent. More generally, controlling for other species in the network will often be important for obtaining reliable estimates of direct (conditional, or all-else-equal) effects. This kind of precision is difficult to obtain from null models, which only test the most extreme possible hypothesis: that *all* direct and indirect interactions are exactly zero. Nevertheless, null models have dominated this field for more than three decades (Strong et al. 1984, Gotelli and Ulrich 2009).

Following Azaele et al. (2010), this paper shows that Markov networks (undirected graphical models also known as Markov random fields; Murphy 2012) can provide a framework for understanding the landscape-level consequences of pairwise species interactions, and for estimating them from observed presence-absence matrices. Markov networks have been used in many scientific fields in similar contexts for decades, from physics (where nearby particles interact magnetically; Cipra 1987) to spatial statistics (where adjacent grid cells have correlated values; Harris 1974, Gelfand et al. 2005). While community ecologists explored some related approaches in the 1980’s (Whittam and Siegel-Causey 1981), they used severe approximations that led to unintelligible results (e.g. “probabilities” greater than one; Gilpin and Diamond 1982).

Below, I demonstrate Markov networks’ ability to produce exact predictions about the direct and indirect consequences of an interaction matrix, and also to make inferences about the species interactions that contributed to an observed set of co-occurrences. Using simulated data sets where the “true” interactions are known, I compare this approach with several existing methods. Finally, I discuss opportunities for extending the approach presented here to other problems in community ecology, e.g. quantifying the overall effect of species interactions on occurrence rates (Roughgarden 1983) and disentangling the effects of biotic versus abiotic interactions on species composition (Pollock et al. 2014).

## Methods

**Markov networks.** Markov networks provide a framework for translating back and forth between the conditional (all-else-equal) relationships among species (Figure 1C) and the kinds of species assemblages that these relationships produce. Here, I show how a set of conditional relationships can determine species composition. Methods for estimating conditional relationships from data are discussed in the next section.

A Markov network defines the relative probability of observing a given vector of species-level presences (1s) and absences (0s), 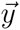 at a site, as

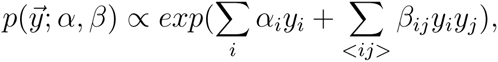

where the second sum is over all 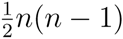 pairs of *n* species. In this model, *α_i_* is an intercept term determining the amount that the presence of species *i* contributes to the log-probability of *y;* it directly controls the prevalence of species *i*. Similarly, *β_ij_* is the amount that the co-occurrence of species *i* and species *j* contributes to the log-probability; it controls the conditional relationship between two species, i.e. the probability that they will be found together, after controlling for the other species in the network (Figure 2A, Figure 2B). For example, if *β_ij_* = +2, then each species’ odds of occurrence would be *e*^2^ times higher when the other one is present (as compared with otherwise equivalent sites). The relative probability of a presence-absence vector increases when positively-associated species co-occur and decreases when negatively-associated species co-occur. As a result, the model tends—all else equal—to produce assemblages where many positively-associated species pairs co-occur and few negatively-associated pairs do (just as an ecologist might expect). When all else is *not* equal (e.g. Figure 1, where the presence of one competitor is associated with release from another competitor), then predicting species’ overall co-occurrence rates can be more complicated, and may require summing over the different possible assemblages (Figure 2B).

**Figure 2:**
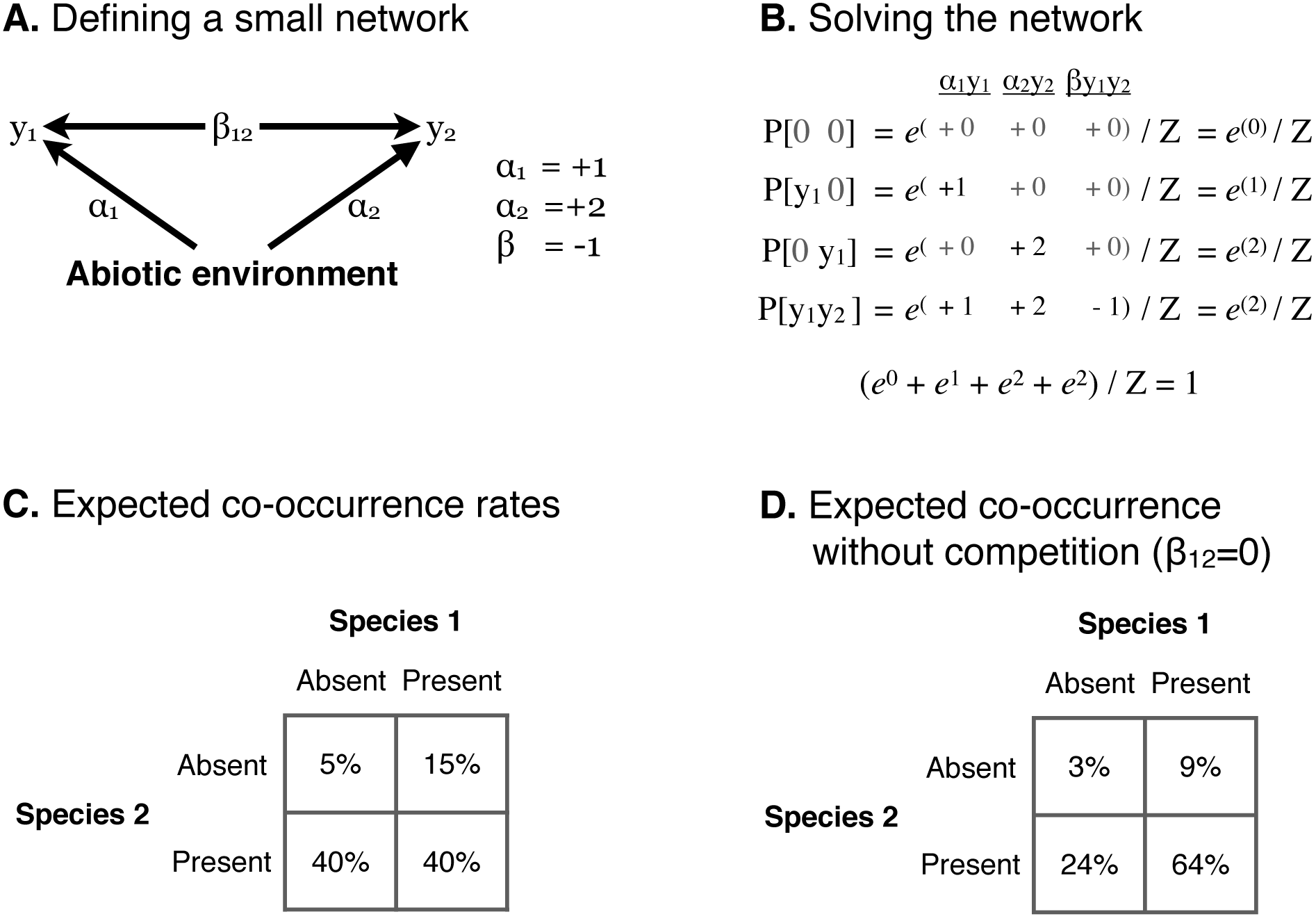
**A.** A small Markov network, defined by its *α* and *β* values. The abiotic environment favors the occurrence of each species (*α* > 0), particularly species 2 (*α*_2_ > *α*_1_). The negative *β*_12_ coefficient is consistent with competition between the two species. **B.** The coefficients determine the probabilities of all four possible presence-absence combinations for Species 1 and Species 2. *α*_1_ is added to the exponent whenever Species 1 is present (*y*_1_ = 1), but not when it is absent (*y*_1_ = 0). Similarly, the exponent includes *α*_2_ only when species 2 is present (*y_2_ =* 1), and includes *β*_12_ only when both are present (*y*_1_*y*_2_ = 1). The normalizing constant *Z*, ensures that the four probabilities sum to 1. In this case, *Z* is about 18.5. **C.** The expected frequencies of all possible co-occurrence patterns between the two species of interest, as calculated in the previous panel. **D.** Without competition (i.e. with *β*_12_ = 0, each species would occur more often.

**Estimating** *α* **and** *β* **coefficients from presence-absence data.** In the previous section, the values of *a* and *β* were known and the goal was to make predictions about possible species assemblages. In practice, however, ecologists will often need to estimate the parameters from an observed co-occurrence matrix (i.e. from a set of independent 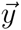 vectors indicating which species are present at each site on the landscape). When the number of species is reasonably small (less than about 30), one can find exact maximum likelihood estimates for all of the *α* and *β* coefficients given a presence-absence matrix by numerically optimizing 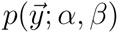. Fully-observed Markov networks like the ones considered here have unimodal likelihood surfaces (Murphy 2012), ensuring that this procedure will converge on the global maximum. I used the rosalia package (Harris 2015a) for the R programming language (R Core Team 2015) to calculate 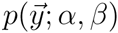 and its gradients (Murphy 2012); the package passes these functions to the “BFGS” method in R’s general-purpose optimizer, which then estimates the Markov network parameters.

**Simulated landscapes.** I simulated several sets of landscapes using known parameters to evaluate different statistical methods’ performance. The first set of landscapes included the three competing species shown in Figure 1. For each of 1000 replicates, I generated a landscape with 100 sites by sampling from a probability distribution defined by the figure’s interaction coefficients (Appendix 1). Each of the methods described below was then evaluated on its ability to correctly infer that the two shrub species competed with one another, despite their frequent co-occurrence.

I then generated landscapes with up to 20 interacting species at 25, 200, or 1600 sites using three increasingly complex models (50 replicates for each combination of size and model; see Appendix 2 for details). I randomly drew the “true” coefficient magnitudes for each replicate landscape from exponential distributions so that most species pairs interacted negligibly but a few pairs interacted strongly enough that their effects could propagate indirectly to other species in the network.

The first set of 20-species landscapes, like the landscapes with three species, were generated directly from a Markov network to ensure that the model could recover the parameters used to generate the “observed” co-occurrence data. Then, I added two environmental factors that varied from location to location across the simulated landscapes, and simulated a new set of co-occurrence data so that species’ *α* coefficients depended on the local environment. The latter set of simulated landscapes provide an important test of the methods’ ability to distinguish co-occurrence patterns that were generated from pairwise biotic interactions from those that were generated by external forces like abiotic environmental filtering. This task was made especially difficult because—as with most analyses of presence-absence data for co-occurrence patterns—the inference procedure did not have access to any information about the environmental or spatial variables that helped shape the landscape (cf Connor et al. 2013). I generated the final set of landscapes with an abundance-based model that included per-capita interaction rates instead of per-species interaction rates.

**Recovering species interactions from simulated data.** I compared seven techniques for determining the sign and strength of the associations between pairs of species from simulated data (Appendix 3). First, I used the rosalia package (Harris 2015a) to fit Markov network models, as described above. For the analyses with 20 species, a weak regularizer (equivalent to a logistic prior with scale 2) ensured that the estimates were always finite.

I also evaluated six alternative methods: five from the existing literature, plus a novel combination of two of these methods. The first alternative interaction metric was the sample correlation between species’ presence-absence vectors, which summarizes their marginal association. Next, I used partial correlations, which summarize species’ conditional relationships. This approach is common in molecular biology (Friedman et al. 2008), but is rare in ecology (see Albrecht and Gotelli (2001) and Faisal et al. (2010) for two exceptions). In the context of non-Gaussian data, the partial correlation can be thought of as a computationally efficient approximation to the full Markov network model (Loh and Wainwright 2013). Because partial correlations are undefined for landscapes with perfectly-correlated species pairs, I used a regularized estimate provided by the corpcor package’s `pcor.shrink` function with the default settings (Schäfer et al. 2014).

The third alternative, generalized linear models (GLMs), also provide a computationally efficient approximation to the Markov network (Lee and Hastie 2012). Following Faisal et al. (2010), I fit regularized logistic regression models (Gelman et al. 2008) for each species, using the other species on the landscape as predictors. This produced two interaction estimates for each species pair; one for the effect of species *i* on species *j* and one for the reverse. These two coefficients are not identifiable from the data, however (Schmidt and Murphy 2012), so I used their average as an overall measure of the pairwise relationship.

The next method, described in Gotelli and Ulrich (2009), involved simulating new landscapes from a null model that retains the row and column sums of the original matrix (Strong et al. 1984). I used the *Z*-scores computed by the Pairs software described in Gotelli and Ulrich (2009) as my null model-based estimator of species interactions.

The last two estimators used the latent correlation matrix estimated by the BayesComm package (Golding and Harris 2015) in order to evaluate the recent claim that the correlation coefficients estimated by “joint species distribution models” provide an accurate assessment of species’ pairwise interactions (Pollock et al. 2014, see also Harris 2015b). In addition to using the posterior mean correlation (Pollock et al. 2014), I also used the posterior mean *partial* correlation, which should control better for indirect effects.

**Evaluating model performance.** For the simulated landscapes based on Figure 1, I assessed whether each method’s test statistic indicated a positive or negative relationship between the two shrubs (Appendix 1). For the null model (Pairs), I calculated statistical significance using its *Z*-score. For the Markov network, I used the Hessian matrix to generate approximate confidence intervals.

For the larger landscapes, I evaluated the relationship between each method’s estimates and the “true” interaction strengths. To ensure that the different test statistics (e.g. correlations versus *Z* scores) were on a common scale, I rescaled them using linear regression through the origin. I then calculated the proportion of variance explained for different combinations of model type and landscape size (compared with a null model that assumed all interaction strengths to be zero).

## Results

**Three species.** As shown in Figure 1, the marginal relationship between the two shrub species was positive—despite their competition for space at a mechanistic level—due to indirect effects of the dominant tree species. As a result, the correlation between these species was positive in 94% of replicates, and the randomization-based null model falsely reported positive associations 100% of the time. Worse, more than 98% of these false conclusions were statistically significant. The partial correlation and Markov network estimates, on the other hand, each correctly isolated the direct negative interaction between the shrubs from their positive indirect interaction 94% of the time (although the confidence intervals overlapped zero in most replicates).

**Twenty species.** In general, each model’s performance was highest for large landscapes with simple assembly rules and no environmental heterogeneity (Figure 3). Despite some variability across contexts, the rank ordering across methods was very consistent. In particular, the four methods that controlled for indirect effects (the Markov network, the generalized linear models, and the two partial correlation-based methods) always matched or outperformed those that did not. The Markov network consistently performed best of all. As anticipated by Lee and Hastie (2012), generalized linear models closely approximated the Markov network estimates (Figure 4A), especially when the data sets were very large (Figure 3). As reviewed in Gotelli and Ulrich (2009), however, most analyses in this field of ecology involve fewer than 50 sites, and the gap between the Markov network and GLMs is larger in this context. As shown in Appendix 4, the standard errors associated with the estimates in Figure 3 are small (less than 0.01), so the differences among methods should not be attributed to sampling error.

**Figure 3:**
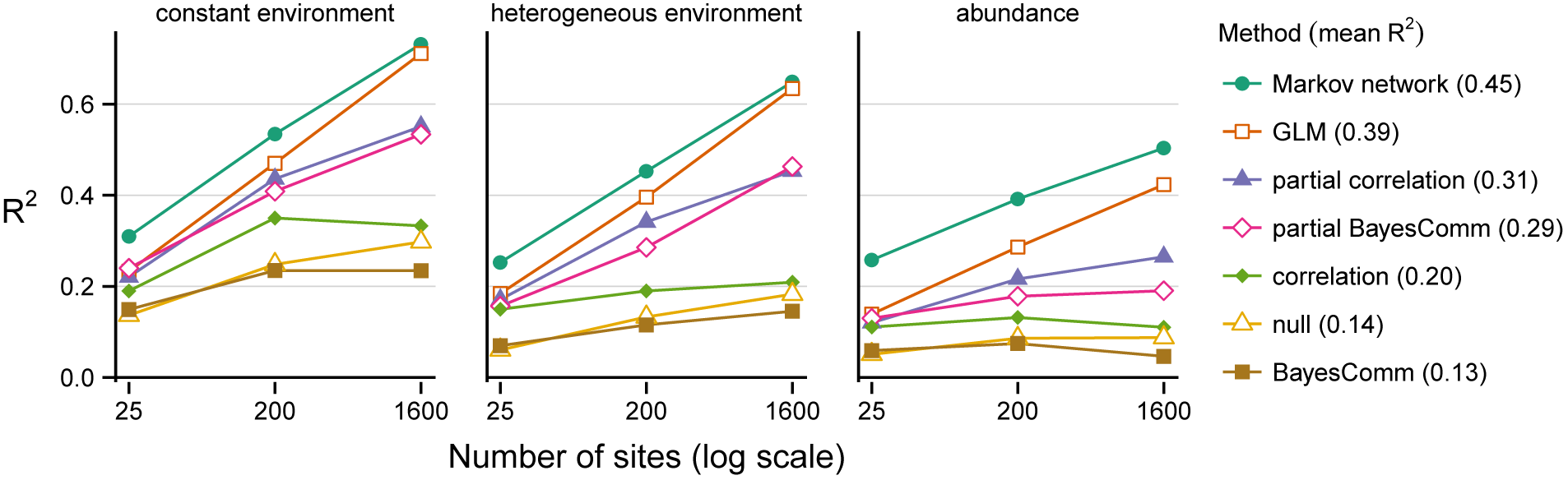
Proportion of variance in interaction coefficients explained by each method versus number of sampled locations across the three simulation types. For the null model (Pairs), two outliers with |*Z*| > 1000 were manually adjusted to |*Z*| = 50 to mitigate their detrimental influence on *R*^2^ (Appendix 5).

Of the methods that did not control for indirect effects, Figure 3 shows that simple correlation coefficients provided a more reliable indicator of species’ true interaction strengths than either the joint species distribution model (BayesComm) or the null model (Pairs). The estimates from these approaches were tightly correlated (after controlling for the size of the landscape) suggesting that the null model only contains a noisy version of the same information that could be obtained more easily and interpretably with simple correlation coefficients (Figure 4B).

**Figure 4:**
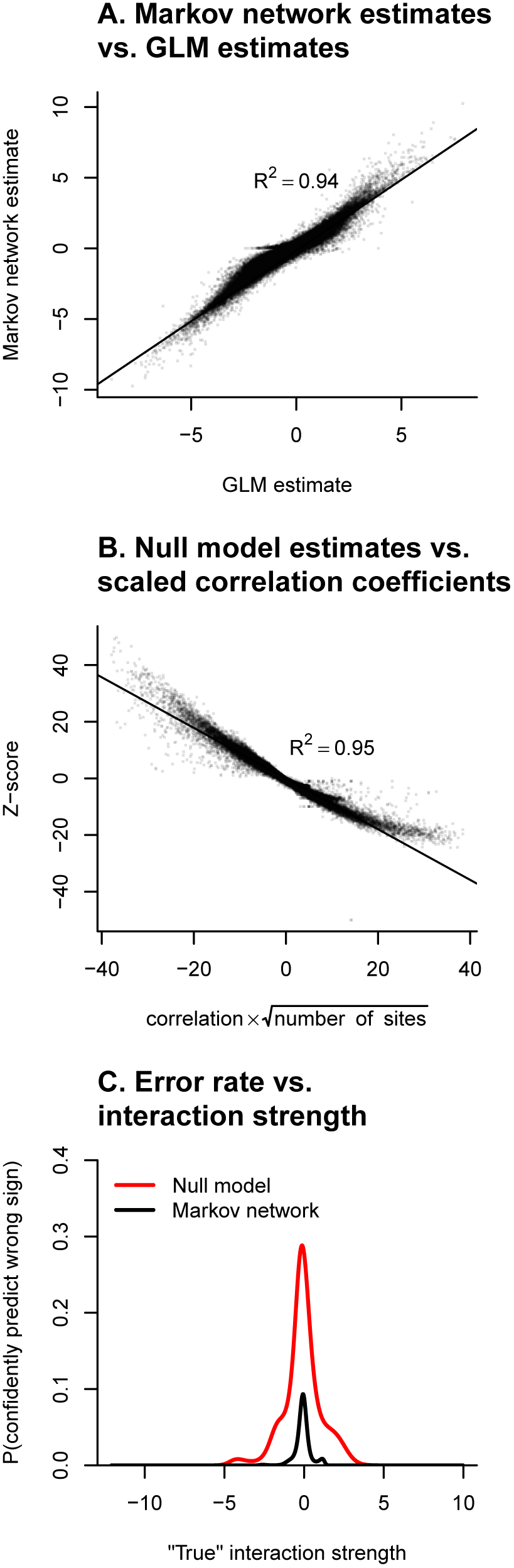
**A.** The Markov network’s estimated interaction coefficients were generally very similar to the GLM estimates. **B.** The null model’s estimates typically matched the (negative) correlation coefficient, after controlling for landscape size. **C.** For any given interaction strength, the null model was much more likely to misclassify its sign with 95% confidence than the Markov network was.

Finally, we can evaluate the models’ statistical inferences (focusing on the first two simulation types, for which the true interaction rates are easiest to interpret). The Markov network’s approximate Type I error rate (defined here as the probability that 0 fell outside the 95% confidence interval for a pair of species where |*β_ij_* | *<* 0.1) depended on the simulation type: 0.02 for simulations that matched the model’s assumptions, versus 0.14 for simulations that included environmental heterogeneity (see Appendix 4 for confidence interval coverage across a range of *β_ij_* values). The null model’s Type I error rates were 0.30 and 0.50 for the constant and heterogeneous landscapes, respectively—far higher than the nominal 0.05 rate. Figure 4C shows, across a range of true interaction strengths, the probability that the null model or the Markov network will predict the wrong sign of the interaction with 95% confidence. The null model makes such errors more than 5 times as often as the Markov network, even though it only reject the null hypothesis twice as often overall (Appendix 4). The Markov network’s errors were also more concentrated around 0, so it never misclassified strong interactions like the null model did.

## Discussion

The results presented above show that Markov networks can reliably recover species’ pairwise interactions from species composition data, even for cases where environmental heterogeneity and indirect interactions cause ecologists’ typical null modeling approaches to reliably fail. Partial covariances and generalized linear models can both provide computationally efficient approximations, but with somewhat lower accuracy (especially for typically-sized data sets with small numbers of sites; Gotelli and Ulrich 2009). The difference in accuracy may be larger for real data sets than for the simulated landscapes in Figure 3, however; linear approximations to the Markov network make larger errors when the interaction matrix is structured (e.g. due to guilds or trophic levels; Loh and Wainwright 2013). Similarly, the separate generalized linear models for each species can severely overfit in some cases (Lee and Hastie 2012). The full Markov network should thus be preferred to the approximations when it is computationally tractable.

Compositional data only contains enough degrees of freedom to estimate one interaction per species pair (Schmidt and Murphy 2012), so none of these methods can identify the exact nature of the pairwise interactions (e.g. which species in a positively-associated pair is facilitating the other). To estimate asymmetric interactions, such as commensalism or predation, ecologists could use time series, behavioral observations, manipulative experiments, or natural history. These other sources of information could also be used to augment the likelihood function with a more informative prior distribution, reducing ecologists’ error and uncertainty relative to Figure 3’s results.

Despite their limitations, Markov networks have enormous potential to improve ecological understanding. In particular, they make many fewer errors than existing approaches, and can make precise statements about the conditions where indirect interactions will overwhelm direct ones. They also provide a simple answer to the question of how competition should affect a species’ overall prevalence, which has important implications for community-level modeling (Strong et al. 1984). Specifically, Equation 1 can be used to calculate the expected prevalence of a species in the absence of biotic influences as *e^α^*/(*e*^0^ + *e^α^*). Competition’s effect on prevalence can then be estimated by comparing this value with the observed prevalence (e.g. comparing Figure 2D with Figure 2C). This novel quantitative result conflicts with most of our null models, which unreasonably assume that prevalence would be the exactly same in the absence of competition as it is in the observed data (Roughgarden 1983).

Markov networks—particularly the Ising model for binary networks—are very well understood, having been studied for nearly a century (Cipra 1987). Tapping into this framework would thus allow ecologists to take advantage of into a vast set of existing discoveries and techniques for dealing with indirect effects, stability, and alternative stable states. Numerous extensions to the basic network are possible as well. For example, the states of the interaction network can be modeled as a function of the local abiotic environment (Lee and Hastie 2012), which would help incorporate networks of biotic interactions into species distribution models (Pollock et al. 2014) and lead to a better understanding of the interplay between biotic and abiotic effects on community structure. Alternatively, models could allow one species to alter the relationship between two other species (Tjelmeland and Besag 1998, cf Bruno et al. 2003).

Finally, the results presented here have important implications for ecologists’ continued use of null models for studying species interactions. Null and neutral models can be useful for clarifying our thinking (Harris et al. 2011, Xiao et al. 2015), but deviations from a particular null model must be interpreted with care (Roughgarden 1983). Even in small networks with three species, it may simply not be possible to implicate specific ecological processes like competition by rejecting a general-purpose null (Gotelli and Ulrich 2009), especially when the test statistic is effectively just a correlation coefficient (Figure 4B). When the non-null backdrop is not controlled for, Type I error rates can skyrocket, the apparent sign of the interaction can change, and null models can routinely produce misleading inferences (Figure 1, Figure 4C, Gotelli and Ulrich (2009)).

Controlling for indirect effects via simultaneous estimation of multiple ecological parameters seems like a much more promising approach: to the extent that the models’ relative performance on real data sets is similar to the range of results shown in Figure 3, scientists in this field could often triple their explanatory power by switching from null models to Markov networks (or increase it nearly as much with linear or generalized linear approximations). Regardless of the methods ecologists ultimately choose, controlling for indirect effects could clearly improve our understanding of species’ direct effects on one another and on community structure.

## Acknowledgements

This work benefited greatly from discussions with A. Sih, M. L. Baskett, R. McElreath, R. J. Hijmans, A. C. Perry, and C. S. Tysor. Additionally, A. K. Barner, E. Baldridge, E. P. White, D. Li, D. L. Miller, N. Golding, N. J. Gotelli, C. F. Dormann, and two anonymous reviewers provided very helpful feedback on the text. This research was partially supported by a Graduate Research Fellowship from the US National Science Foundation and by the Gordon and Betty Moore Foundation’s Data-Driven Discovery Initiative through Grant GBMF4563 to E. P. White.

